# goCTF: Geometrically optimized CTF determination for single-particle cryo-EM

**DOI:** 10.1101/426189

**Authors:** Min Su

## Abstract

Preferred particle orientation represents a recurring problem in single-particle cryogenic electron microcopy (cryo-EM). A specimen-independent approach through tilting has been attempted to increase particle orientation coverage, thus minimizing anisotropic three-dimensional (3D) reconstruction. However, focus gradient is a critical issue hindering tilt applications from being a general practice in single-particle cryo-EM. The present study describes a newly developed geometrically optimized approach, goCTF, to reliably determine the global focus gradient. A novel strategy of determining contrast transfer function (CTF) parameters from a sector of the signal preserved power spectrum is applied to increase reliability. Subsequently, per-particle based local focus refinement is conducted in an iterative manner to further improve the defocus accuracy. Novel diagnosis methods using a standard deviation defocus plot and goodness fitting heatmap have also been proposed to evaluate CTF fitting quality prior to 3D refinement. In a benchmark study, goCTF processed a published single-particle cryo-EM dataset for influenza hemagglutinin trimer collected at a 40-degree specimen tilt. The resulting 3D reconstruction map was improved from 4.1Å to 3.7Å resolution. The goCTF program is built on the open-source code of CTFFIND4, which adopts a consistent user interface ease of use.

## 1. Introduction

Recent advances in cryo-EM emanate from modern technological innovations. One of these major advancements is the improvement of the electron microscope design, allowing a vitrified specimen to sit on an ultra-stable compustage for multiple days, which is suitable for collecting large datasets through automation. This newly designed compustage has benefitted cryogenic electron tomography (cryo-ET) and, along with the improved image processing technique (Mattei et al., 2016; Schur et al., 2016), has advanced reconstructed tomograms toward near-atomic resolution. Single-particle cryo-EM, in contrast, has not taken full advantage of this compustage innovation, particularly for image acquisition at specimen tilt. Preferred particle orientation represents a common, recurring problem for cryo-EM (Glaeser, 2016; Glaeser and Han, 2017), which often causes anisotropic map resolution. In extreme cases, the particle alignments could be completely wrong due to missing views, thus misrepresenting the true structure of an object.

Various approaches have been explored to solve preferred particle orientation (Chowdhury et al., 2015; Frank et al., 1991; Liu et al., 2013; Lyumkis et al., 2013; Meyerson et al., 2014; Nguyen et al., 2015); however, these methods are only occasionally effective in increasing the range of particle orientation. Additionally, these approaches are highly specimen-dependent and far from reproducible. Recent efforts have attempted to address preferred specimen orientation in single-particle cryo-EM through tilting, which is a specimen-independent and general approach. A previous study (Su et al., 2017) showed that AAA ATPase Vps4 oligomers have a highly preferred top-down orientation in ice. By including tilted images, the open and closed state oligomers could be differentiated at the near-orthogonal view. In another study (Tan et al., 2017), the authors demonstrated for the first time that single-particle cryo-EM is capable of reaching near-atomic resolution with the data collected at a 40-degree specimen tilt, using influenza hemagglutinin trimer as a benchmark sample.

These two recent examples from different groups showed the benefits afforded to single-particle cryo-EM through tilting, even though this strategy is generally considered challenging and thus impractical for various technical limitations in cryo-EM field. One challenge associated with tilting is that a geometry-induced thicker specimen tends to accumulate more electron charging, thus enhancing the phenomenon of beam-induced specimen movement as local events in addition to global drift, which deteriorates the motion correction performance. The suboptimal drift correction decreases the imaging quality with a degraded signal-to-noise ratio (SNR), which often is considered unsuitable for high-resolution single-particle cryo-EM.

Another known limitation is the focus gradient at specimen tilt, particularly because there is no widely adopted general approach to tackle the CTF gradient for single-particle cryo-EM tilting. Attempts have been made, with some success, to use the popular software gCTF to address the focus gradient by taking advantage of the software’s local focus refinement option (Zhang, 2016). However, gCTF was not specifically designed for determining the focus gradient with its local refinement strategy; thus, its performance is suboptimal in practice. Moreover, the gCTF focus refinement strategy is sensitive to local region SNR in a cryo-EM micrograph, which is a common issue associated with tilting. In addition, the refined focus accuracy in gCTF is highly reliant on particle abundance at the local region of interest (ROI).

In the present study, goCTF, a new program explicitly designed to determine the focus gradient in single-particle cryo-EM, is implemented and validated using a novel geometrically optimized approach. The goCTF program adopts an ultra-reliable strategy in CTF determination, suitable for noisy and drifting cryo images taken at high specimen tilt. This new software tool was developed based upon the open-source code of CTFFIND4 (Rohou and Grigorieff, 2015); hence, it inherited the same input parameter convention for easy application. The goCTF program as an alternative for CTF determination and local focus refinement can be plugged into the user interface of the software RELION (Scheres, 2012) and can easily be inter-converted to other software.

## 2. Theory and methods

### 2.1 Definition of contrast transfer function

The CTF of an electron microscope can be described using the following equation (Mindell and Grigorieff, 2003):

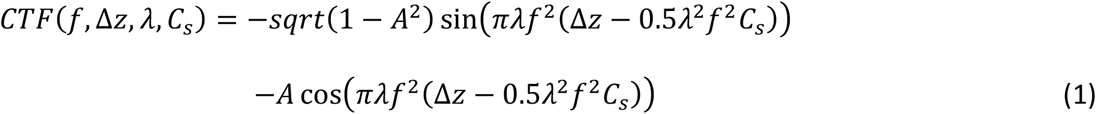

In this equation, *f* denotes the spatial frequency, Δ*z* the defocus, *λ* the electron wavelength, *C_s_* the spherical aberration of the microscope, and *A* the relative amplitude contrast.

### 2.2 Focus gradient impact

The micrograph collected at specimen tilt is modulated by the CTF with a focus gradient perpendicular to the stage tilt axis. To quantitatively evaluate the focus gradient effect across the micrograph, one representative simulation is conducted at a tilt angle of 45 degrees to illustrate this impact as a function of resolution in reciprocal space. In the simulation, the following microscope parameters are adopted: HT 300 keV, Cs 2.7, amplitude contrast 0.1 and pixel size 1.0. These parameters closely represent a Titan Krios. Circular boxes (256 pixels in diameter) are placed on the micrograph, resulting in a relative focus difference of 256Å between neighboring colored columns of boxes (Fig. 1a and b). The reference black column of boxes is set to be at 2.0 μm defocus, with the corresponding paired CTF curves (black vs. blue, black vs. green and black vs. orange) plotted in Fig. 1c (top, middle and bottom, respectively). This simulation clearly shows that the paired column of boxes located 1024Å (black vs. orange) apart, perpendicular to the tilt axis, exhibit a 90-degree phase shift at 5Å resolution, and these regions undergo the complete opposite phase shift at 4Å resolution. This simulation demonstrates that, in order to pursue a near-atomic resolution reconstruction for single-particle cryo-EM through tilting, it is essential to take the focus gradient into account for CTF correction.

**Fig. 1.**
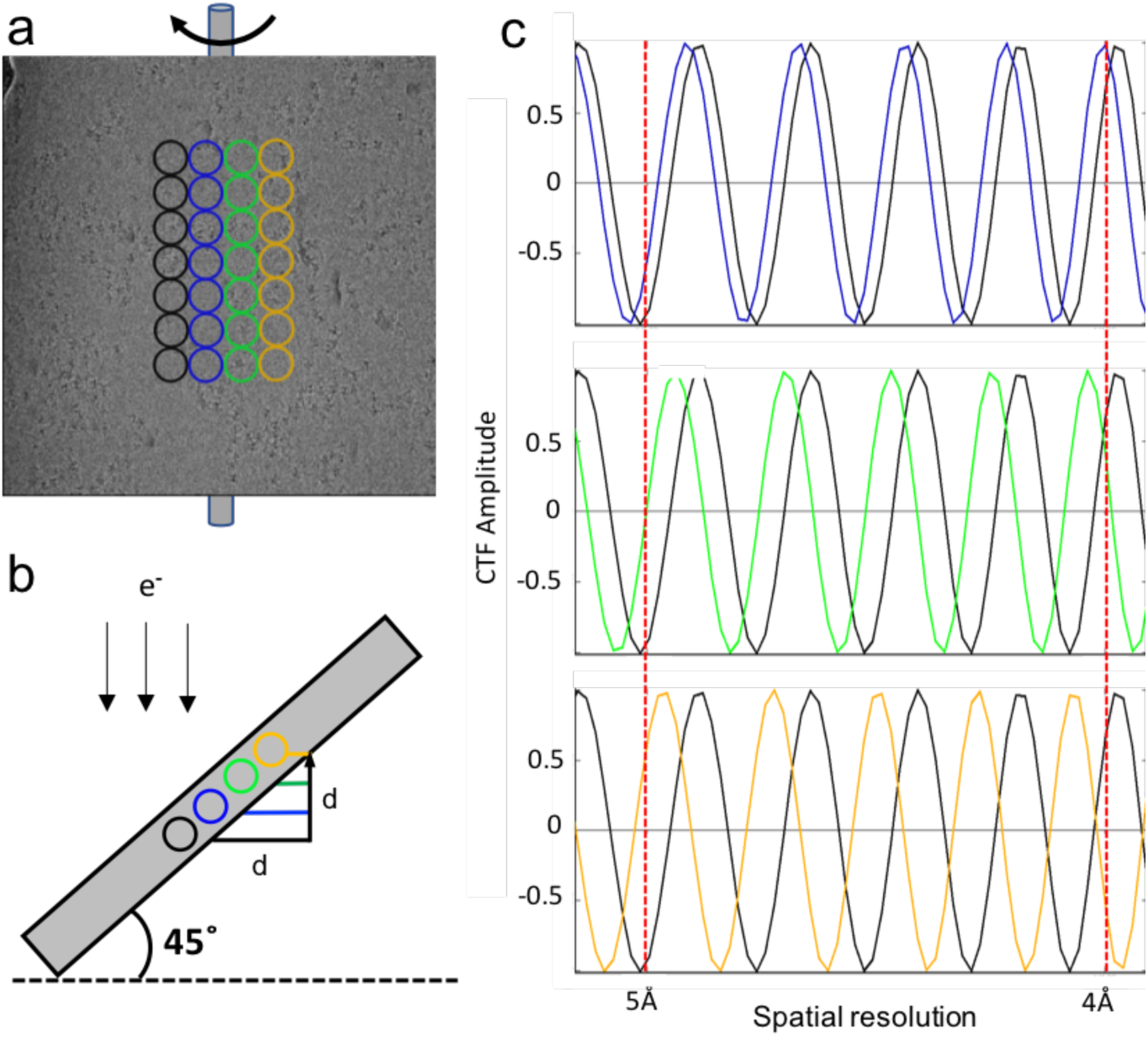
CTF focus gradient analysis perpendicular to the tilt axis. (a) One representative cryo-EM micrograph collected at a 45-degree specimen tilt. Each column of colored circular boxes is placed parallel to the tilt axis with a box size of 256Å in diameter. (b) Schematic showing projected side view indicating the focus gradient across the micrograph. (c) Simulated CTF curves plotted in pairs (top: black vs. blue; middle: black vs. green; bottom: black vs. orange) indicating relative phase shift due to the focus gradient of each pair respectively.

### 2.3 Determination of defocus from anisotropic power spectrum

Beam-induced specimen movement is a severe problem, particularly for imaging while tilted in cryo-EM, with specimen movement present both as global and as local drift on the micrograph. Although gold grids have been proposed to minimize drift for tilt application (Russo and Passmore, 2014), drift is generally unpredictable and highly specimen dependent. Thus, it is commonly observed that the power spectrum from a tilted micrograph appears to be missing information directionally, with a stronger spectrum and a diminished spectrum orientated orthogonally (Fig. 2a).

**Fig. 2.**
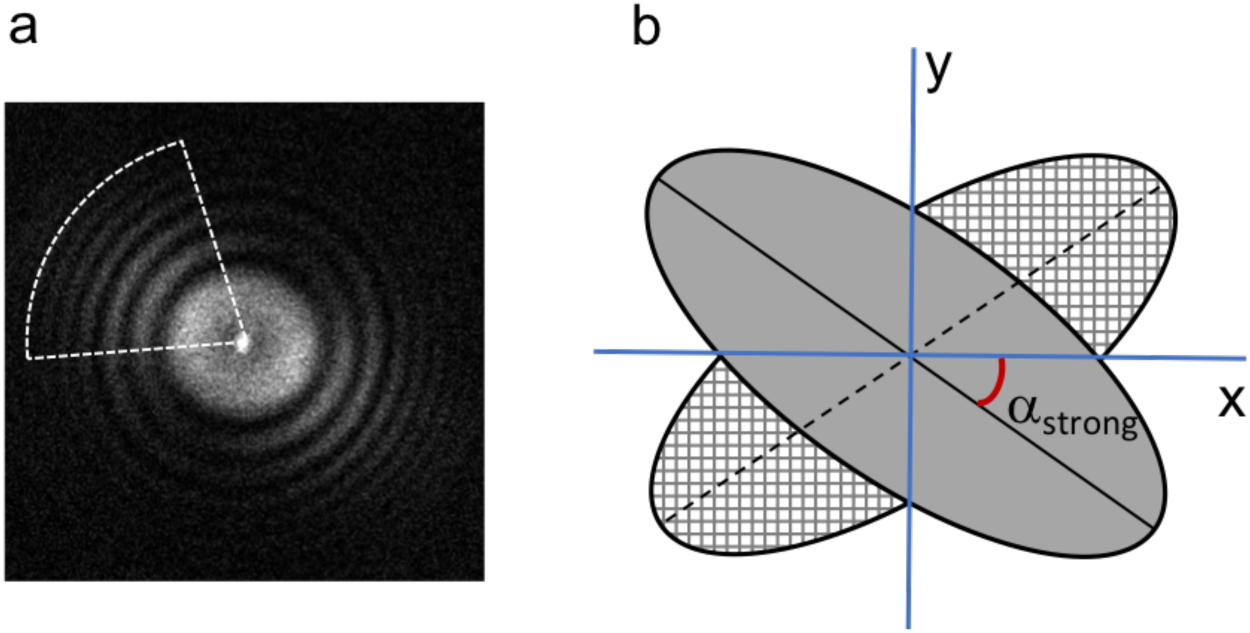
CTF determination from an anisotropic power spectrum. (a) Typical power spectrum from a tilted cryo-EM micrograph indicating anisotropic signal scattering. The white color sector indicates the region that preserves the most signal useful for CTF estimation. (b) Diagram showing the preserved power spectrum in a gray ellipse and its mirror along the x axis.

To my knowledge, most software programs compute CTF parameters using the full range of a power spectrum, regardless of its anisotropic resolution (Ludtke et al., 1999; Mallick et al., 2005; Mindell and Grigorieff, 2003; Penczek et al., 2014; Rohou and Grigorieff, 2015; Shaikh et al., 2008; Sorzano et al., 2004; Tang et al., 2007; Vargas et al., 2013; Voortman et al., 2011). The conventional search is either conducted in two dimensions (2D) to determine stigmatism or reduced in one dimension (1D) to a detrimental rotationally averaged power spectrum. In reality, as a result of the geometric defect of tilting, micrographs display an anisotropic power spectrum particularly from high tilt. In combination with local drift events, it remains challenging to decouple astigmatism vs. anisotropic resolution from a power spectrum due to the degraded SNR at tilt. Moreover, it is generally observed that beam induced image drift is one of the dominant factors leading to decreased image quality for tilt applications in cryo-EM. Therefore, it is more practical and reliable to compute CTF parameters within a confined sector along the orientation of the stronger power spectrum, instead of using the full range of the power spectrum by rotational average.

To estimate the orientation of the stronger power spectrum, goCTF adopted an algorithm (van Heel et al., 2000) that was originally developed to determine the astigmatism angle from a micrograph and later implemented in CTFFIND4. First, the preprocessed power spectrum is mirrored along one of its axes (Fig. 2b). It is then aligned rotationally against its mirrored spectrum using a 1D exhaustive search by maximizing the correlation efficiency. In the case of tilting, missing directional information is a dominant effect; thus, this algorithm essentially estimates the anisotropic resolution of a power spectrum, rather than astigmatism.

### 2.4 Local focus refinement strategy

Although single-particle cryo-EM through tilting has shown great potential toward achieving near-atomic resolution reconstructions, it generally remains challenging, particularly in high-quality data collection, due to severe beam induced image drift, which is difficult to completely correct by drift correction software. Subsequently, the local focus refinement strategy in the known program, gCTF, often produces suboptimal results when applied to lower-quality images. In the extreme case, gCTF could completely fail due to low SNR in a local ROI within a tilted micrograph. A novel, reliable approach is essential to fulfill per-particle focus refinement in the case of tilt for single-particle cryo-EM.

Focus gradient is one of the critical issues preventing single-particle tilt application from being a general practice in the cryo-EM field. Due to an anisotropic power spectrum and extremely low SNR at a given local ROI in a tilted cryo-EM micrograph, it remains challenging and inaccurate to determine the stage tilt angle coupled with the tilt axis orientation simultaneously. The goCTF program assumes that the detector pixel array has been assembled and aligned approximately colinearly with the stage tilt axis by the manufacturer. Thus, one revised strategy of searching for the global focus gradient can be simplified to model the stage geometry at a given tilt angle with no discernable effect.

To determine the focus gradient, the newly developed program goCTF follows the workflow shown in Fig. 3c. In the first step, a tilted micrograph is divided into parallel tiles along the tilt axis. The defocus value is computed *ab initio* from each individual tile, then fitted by a linear regression model to define the global focus gradient distribution across the micrograph (Fig. 3a and b). In the next step, the defocus value of a particle of interest at any given location is extrapolated according to the modeled focus gradient distribution. This extrapolated defocus serves as the starting value for the next round of iterative refinement within a constrained local focus search range and an adaptive resolution search range in reciprocal space. In the first iteration of local refinement, a larger rectangular area (cyan) surrounding the particle of interest (green circle) is used to compute the power spectrum to minimize anisotropic spectrum defect resulting from stage tilt. The refined local defocus value from the first iteration is then subjected to a second round of refinement in the same manner, but using a smaller cropped area (orange) to more closely represent the true focus value of a particle of interest. A modified scoring function for each iteration, *CC_i_*, is then applied to calculate the normalized cross-correlation coefficient between the modeled CTF and amplitude spectrum *A_di_* as a target function for local CTF refinement:

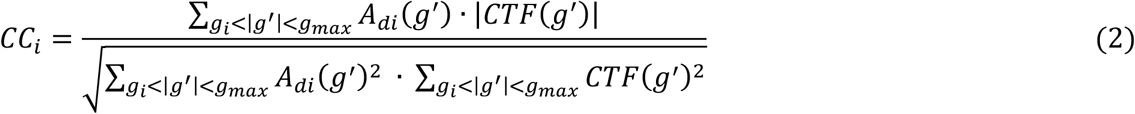

**Fig. 3.**
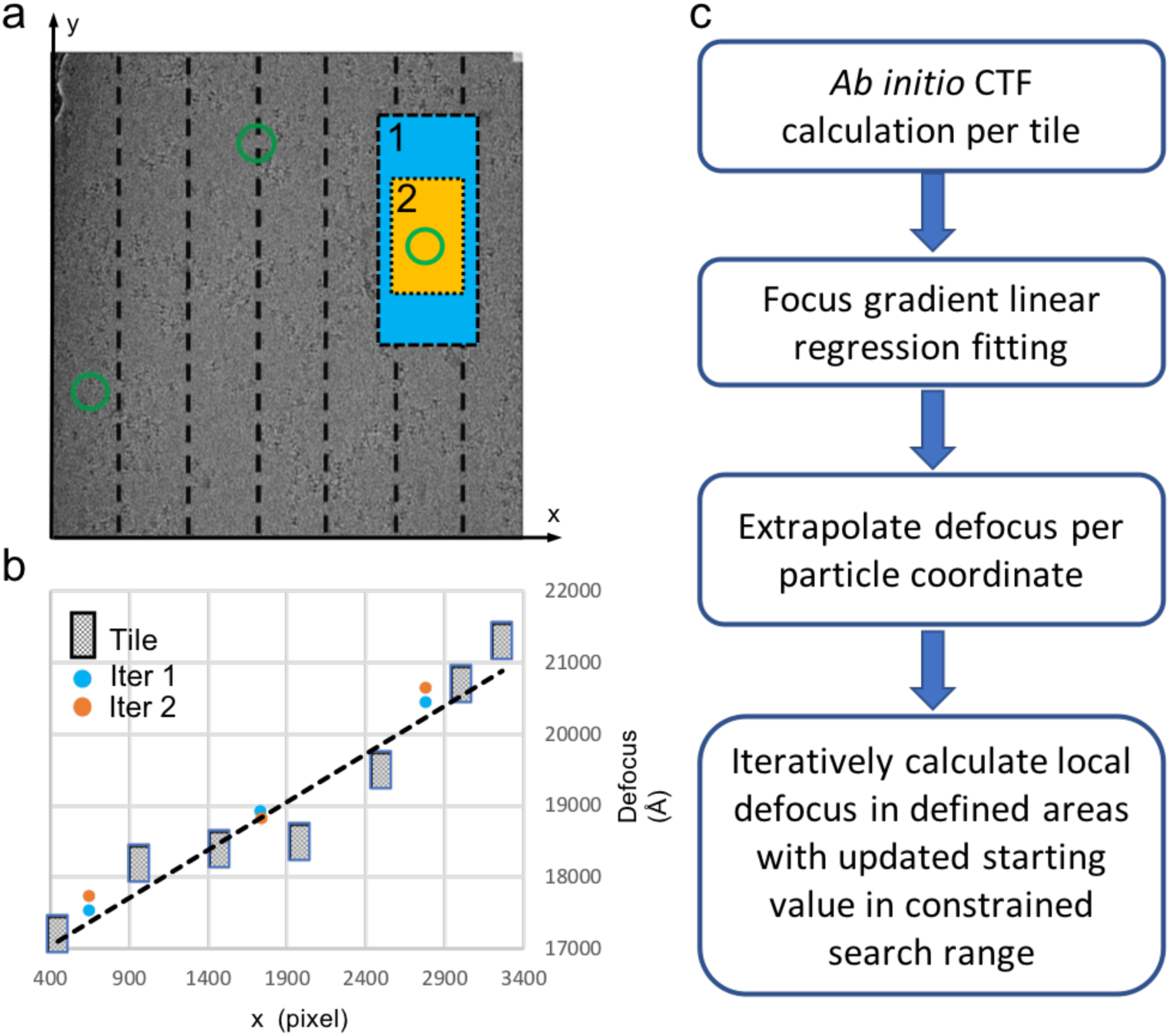
Geometrically optimized local focus refinement strategy. (a) One representative cryo-EM micrograph is divided by dashed lines into tiles parallel to the tilt axis. Green circles represent selected particle coordinates within a micrograph, with one particle of interest cropped by a larger rectangle box (cyan) and smaller rectangle box (orange) for local CTF refinement in iteration 1 and 2 iteratively. (b) The defocus value is calculated from each individual tile and subsequently used for global focus gradient estimation modeled by a linear regression fitting shown with the dashed line. The local focus refinement results are indicated by cyan and orange dots for iteration 1 and 2 respectively. (c) Flow chart of goCTF.

where *g*′ denotes that the spatial frequency lies in the pre-defined sector of strong amplitude spectrum in *A_di_*.

### 2.5 Local focus refinement quality analysis

It is challenging to accurately determine the local focus from an experimental tilted cryo-EM micrograph; therefore, it is essential to analyze the performance of this geometrically optimized strategy for high-resolution reconstruction as a reliable off-the-shelf tool. Grid boxes were placed across the entire micrograph, with each column labeled with alternating blue or red dots. The 3D and 2D plots are displayed in Fig. 4a and b, respectively, showing the global focus gradient consistent with the known stage tilt angle of 45 degrees. The refined local foci appear to deviate from the extrapolated values, perhaps indicating particle defocus change within bulk ice thickness. The 3D plot, together with 2D plot, are undoubtedly important and intuitive for validating the estimated global focus gradient.

**Fig. 4.**
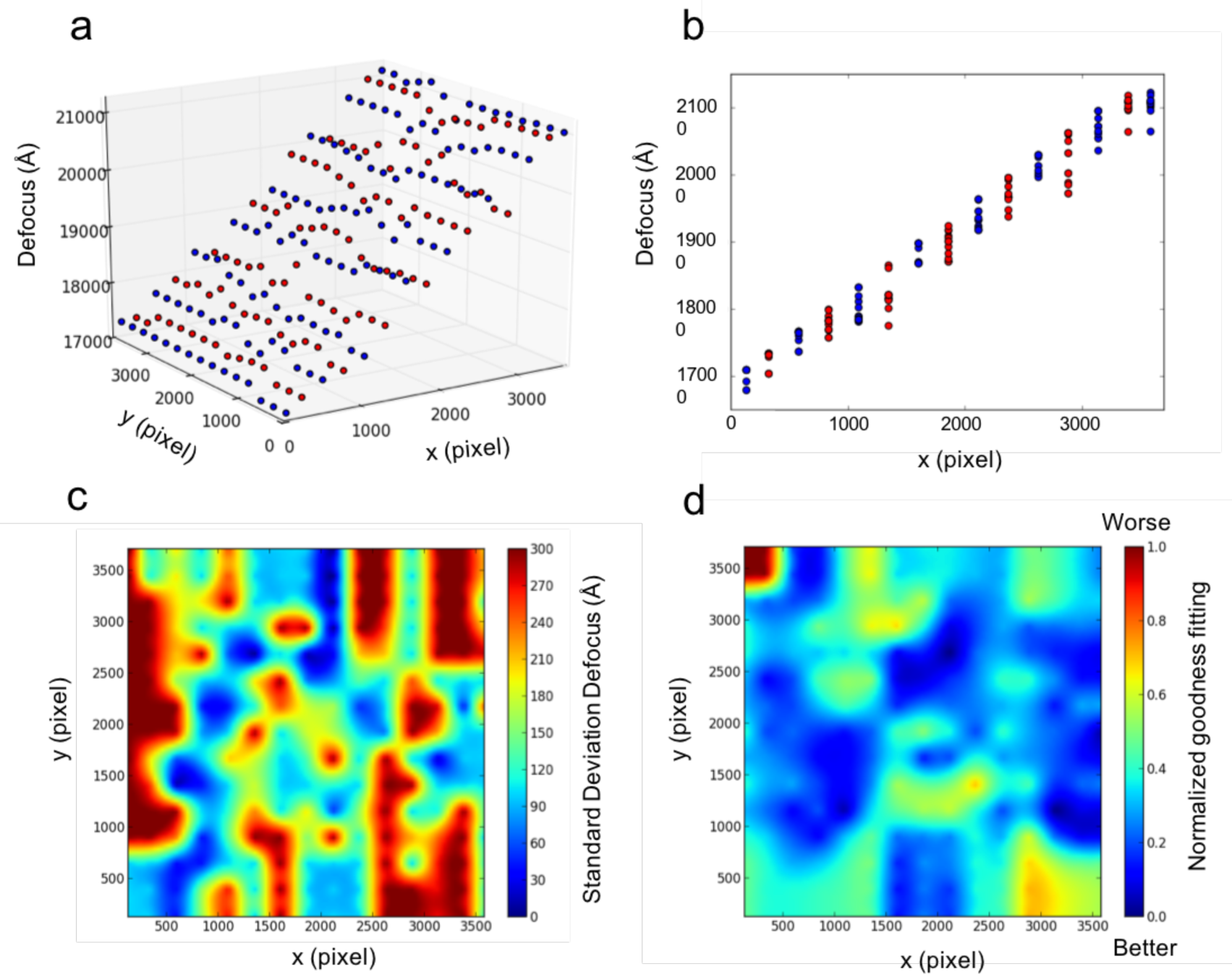
Local focus refinement analysis. (a) Grid boxes were placed across the micrograph in Fig. 3a with each column labeled by alternating blue or red dots. The refined local focus gradient is plotted in 3D. (b) The focus gradient is projected along the y axis and plotted in 2D. (c) The standard deviation defocus map indicates refined local defocus deviation from the fitted global focus gradient. (d) The normalized goodness fitting of local refinement is plotted in the heatmap, indicating agreement between the simulated CTF model and experimental power spectrum.

To further analyze the local focus dynamic behavior with the freedom deviating from the modeled global gradient during the iterative refinement process, the square root of the difference between the extrapolated value from the global gradient and final refined local value, called the standard deviation (S.D.) defocus map, is plotted in Fig. 4c. This S.D. map is informative for indicating local focus variations but also could be adopted to evaluate ice thickness across a micrograph.

Some local focus variation from a global gradient in a titled micrograph is expected, whereas little confidence remains without a CTF fitting quality measurement. Here, goCTF adopts a similar strategy as implemented in CTFFIND4 to measure the correlation between the fitted CTF model and the calculated experimental power spectrum for each individual local box. This measured goodness fit is quantified in a unit of Å, then normalized by the range between the largest and smallest value of the goodness of fit among all the particles within a micrograph. This normalized relative goodness of fit for the local defocus is plotted in a heatmap in Fig. 4d, which indicates the quality of fitting across the entire micrograph and could also be translated to an evaluation of micrograph at a local given ROI.

## 3. Results

### 3.1 Validation with graphene oxide grid

Because simulating a micrograph modulated by CTF with a focus gradient is not a straightforward process, an alternative approach is explored using a commercial graphene oxide grid to validate goCTF performance in the case of weak signal scattering. Graphene oxide films are extremely thin and frequently used as the support substrate in cryo-EM studies for challenging biological macromolecules (Pantelic et al., 2011). The monolayer of graphene oxide generates minimal background noise in low-dose conditions under electron microscope; thus, it serves as an ideal material to test the reliability of goCTF with low signal scattering mimicking a typical cryo-EM micrograph.

Test images were collected using a Tecnai 12 operated at 120 keV with an Ultra4000 CCD camera at a total dose ~ 40 e/A^2^. A representative image of graphene oxide is shown in Fig. 5a, with its diffraction pattern displayed on the bottom right indicating a monolayer of graphene.

**Fig. 5.**
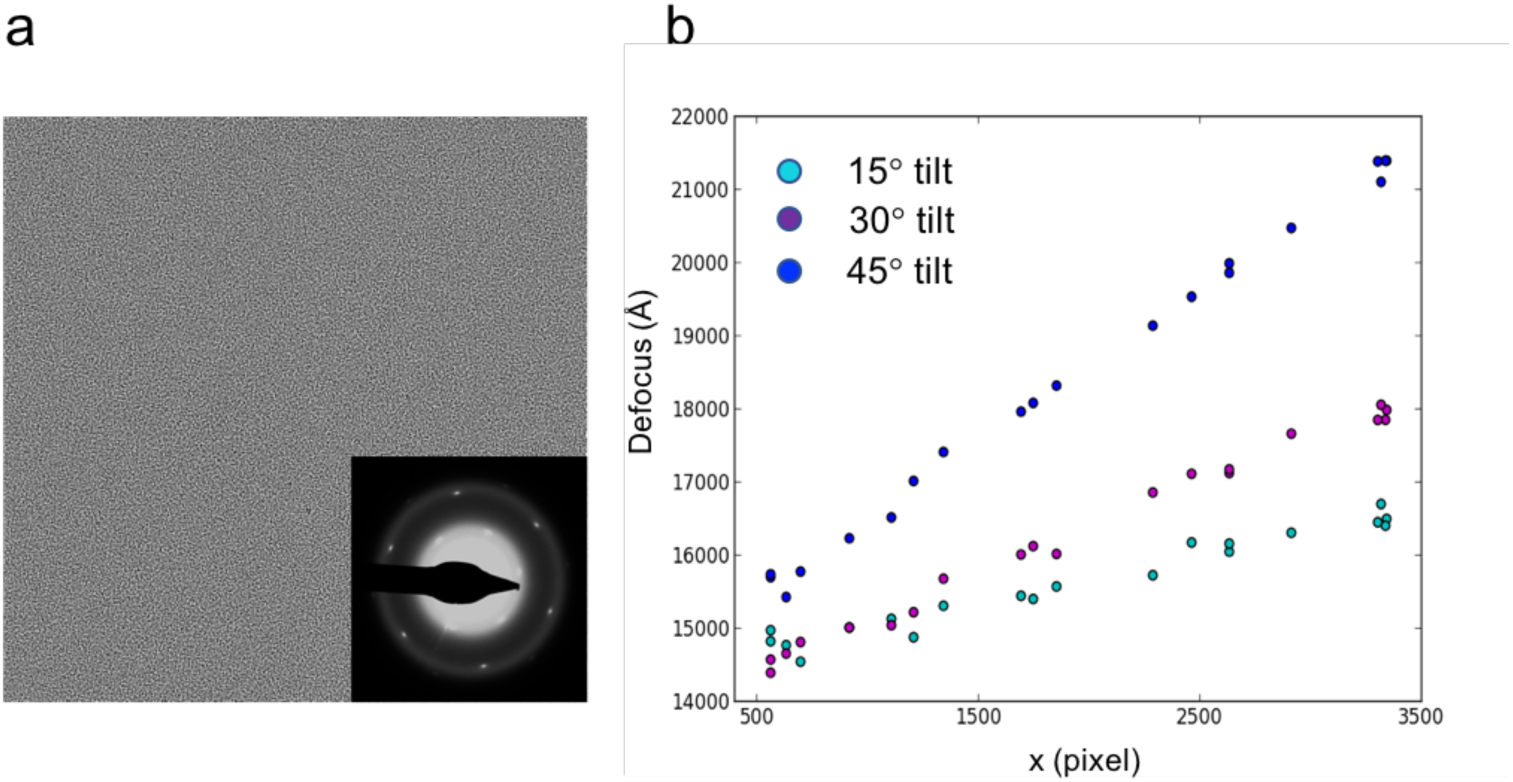
Validation with a graphene oxide grid. (a) One untilted image is taken from the graphene oxide film with its diffraction pattern shown on the bottom right, indicating a monolayer of graphene substrate. (b) Tilt images were collected at 15, 30 and 45 degrees with 20 boxes randomly placed across the corresponding micrograph for local CTF refinement shown in cyan, purple and blue dots, respectively.

Three images were taken at specimen tilt of 15, 30 and 45 degrees, respectively. Twenty boxes with random positions were selected and applied to all three images to test the local focus refinement performance. It is encouraging to observe that goCTF successfully differentiated all three tilt angles in the case of the low signal scattering of graphene oxide (Fig. 4b), suggesting the robustness of this program and its suitability for cryo-EM tilt applications.

### 3.2 Comparative test in accordance with particle density distribution

An experimental cryo-EM image likely contains particles with various density distributions, resulting in varying strengths of signal scattering. It is particularly challenging for local defocus refinement in the case of lower particle density distribution in a given ROI. Therefore, it is essential to conduct reliability tests of goCTF that cover the full range of experimental scenarios. Moreover, it would be more convincing to compare goCTF with a known and proven program in all the tests.

The well-known software gCTF, which has been proven capable of refining local defocus, has been used previously in single-particle cryo-EM through tilt application. Comparative performance tests between gCTF and goCTF were conducted using influenza hemagglutinin (HA) trimer as a benchmark test sample (EMPIAR-10097), since this dataset was imaged at a 40-degree specimen tilt. The focus refinement results from gCTF were obtained from the deposit particle stack file as part of the benchmark dataset and plotted for the following comparisons: Fig. 6a and b (top) show representative cryo-EM micrographs from this test dataset with high and low particle density distributions, respectively; 50, 10 and 5 boxes with random positions were chosen and applied to both images to compare the reliability of goCTF and gCTF in the two test cases with high and low SNR.

**Fig. 6.**
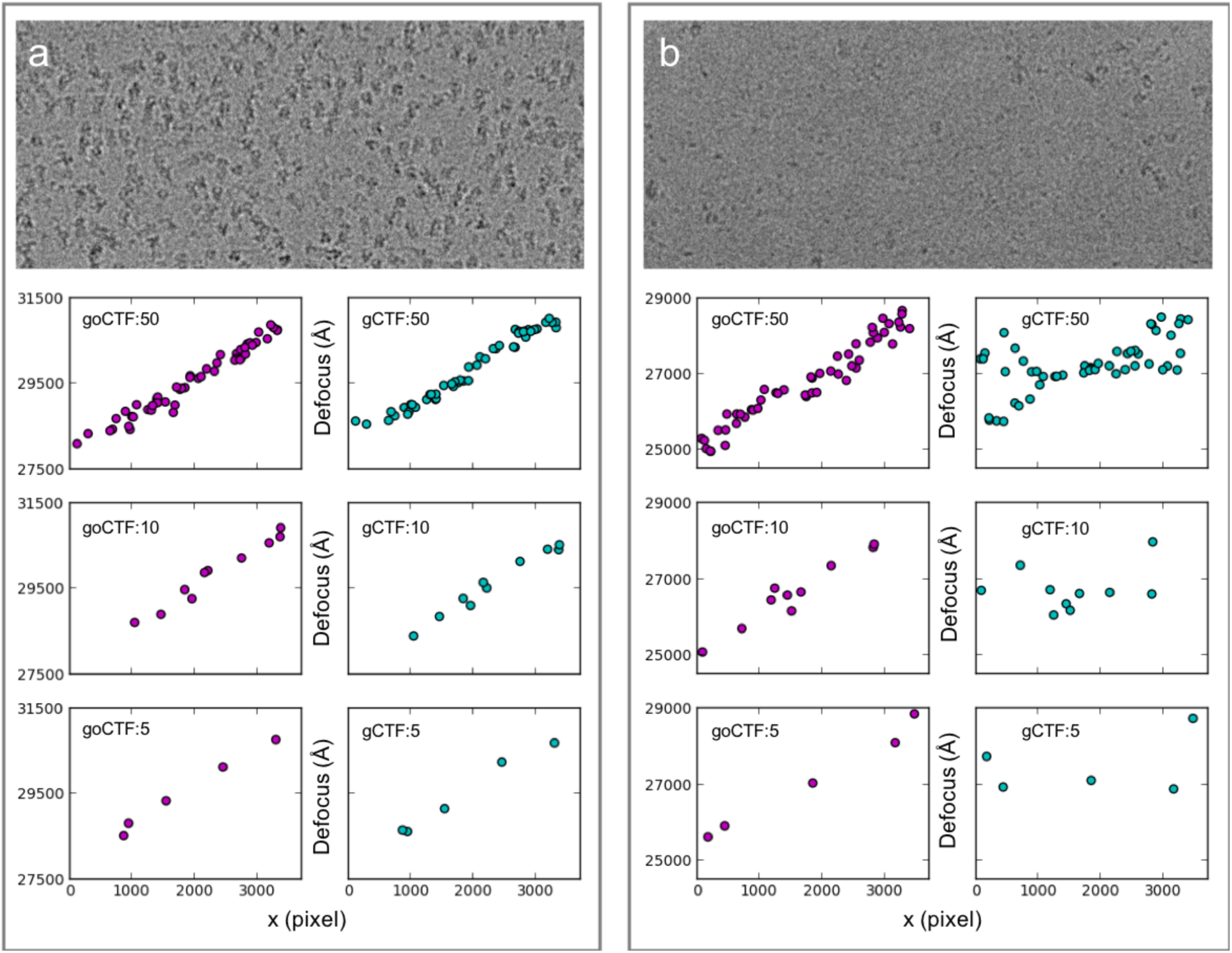
Comparison of the particle abundance impact between goCTF and gCTF. (a) One representative micrograph from a benchmark dataset (EMPIAR-10097) showing high particle abundance (top) and (b) low particle abundance. For both (a) and (b) 50, 10 and 5 boxes with random positions were placed on the micrographs for local focus refinement. The determined focus gradient was plotted side-by-side for comparison between goCTF (purple dots) and gCTF (cyan dots).

In the case of high particle density distributions in Fig. 6a, both goCTF and gCTF show mutually consistent results independent from the number of particle selected. In the case of low particle density distributions in Fig. 6b, in contrast, gCTF local refinement starts diverging near both edges of the micrograph (Fig. 6b, gCTF:50), whereas goCTF demonstrates robust results, clearly indicating a focus gradient representative of a 40-degree tilted micrograph (Fig. 6b, goCTF:50). This discrepancy occurs because gCTF retrieves the global averaged defocus of a micrograph as the initial search value to start local refinement. On the two far ends of a micrograph, this initial value is far from the global average due to tilt. In addition, the low abundance of neighboring particles results in a low averaged power spectrum, thus causing the local refinement to deviate from the true value. When the box number is reduced to 10, gCTF local refinement shows large variations barely indicating an expected focus gradient (Fig. 6b, gCTF:10). In comparison, goCTF exhibits a reasonable focus gradient across the micrograph, testifying that its local refinement strategy is independent from neighboring particles (Fig. 6b, goCTF:10). When the box number is further reduced to only 5, gCTF local refinement fails completely (Fig. 6b, gCTF:5); goCTF, however, remains consistent with the previous results, demonstrating a clear global focus gradient in this extreme case (Fig. 6b, goCTF:5). Further substantial comparison tests using this benchmark dataset are shown in Fig. S1. It is worth mentioning that the gCTF local refinement strategy takes advantage of pre-selected neighboring particles to enhance the averaged power spectrum for CTF fitting. This strategy is beneficial in refining local defocus with no or small angle tilt. In the case of high tilt however, its local refinement reliability and accuracy may be overly influenced by neighboring particle abundance and distributions perpendicular to the tilt axis.

### 3.3 Benchmarking in 3D refinement

As demonstrated in the results shown above, the geometrically optimized approach, goCTF, showed superior reliability in determining the focus gradient in comparison with the proven known program gCTF. In addition to simply determining the focus gradient, goCTF also was tested to determine whether the superior performance of this program could be translated into a better reconstruction map after 3D refinement.

A comparative 3D refinement test was conducted using the published single-particle cryo-EM dataset (EMPIAR-10097), which contains 130,000 particles of influenza hemagglutinin trimer. This dataset was reprocessed following a Homogenous refinement approach using the software cryoSPARC (Punjani et al., 2017). With the original dataset, cryoSPARC reached a global resolution of 4.1Å reconstruction showing an equivalent quality map compared to the reported map using RELION. The same dataset in cryoSPARC, with the defocus updated by goCTF, reported a global resolution of 3.7Å showing noticeably improved map quality (Fig. 7a, Fig. S2). Furthermore, directional 3DFSC plots were generated using server (https://3dfsc.salk.edu), indicating improved FSC histogram and reconstruction isotropy particularly in Z direction (Fig. 7b).

**Fig. 7.**
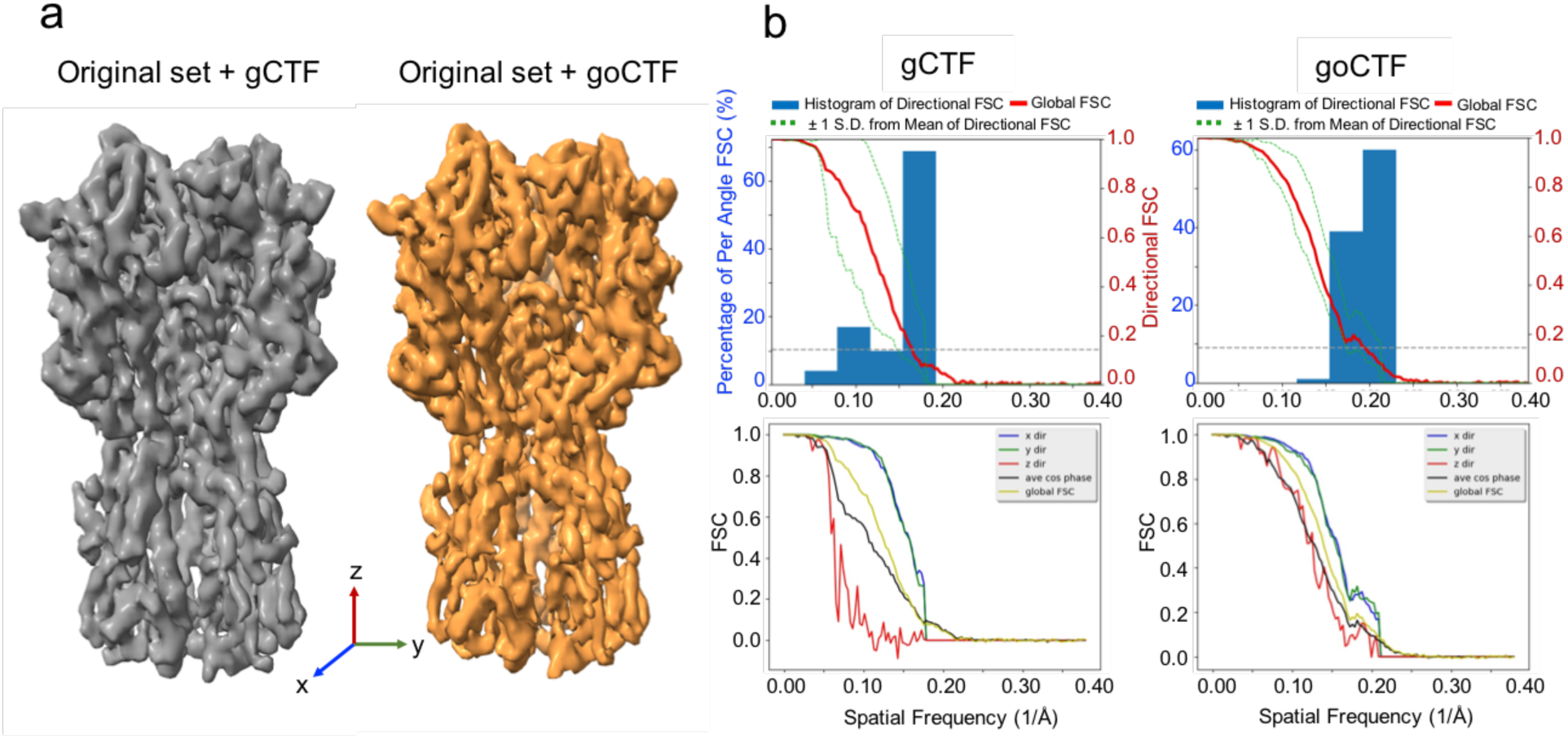
Benchmark in 3D refinement using cryoSPARC with published dataset (EMPIAR-10097) containing influenza hemagglutinin trimer. (a) Isosurface rendering of the 3D map results from the original dataset (130,000 particles) processed by gCTF shown in grey, and the original dataset updated defocus by goCTF shown in orange. (b) Directional 3DFSC plots shown from gCTF (left) and goCTF (right).

## 4. Discussion

The goCTF program was developed to target individual particle focus refinement with an adaptive search range instead of *ab initio*. In addition, local focus can be further refined in an iterative manner to converge on a true defocus at the region of interest. Due to uneven ice thickness in cryo-EM images (Noble et al., 2018), global focus gradient likely differs from a simplified geometry model as a flat plane. Thus, one such coarsely determined gradient might be sufficient to approximate an adaptive search range for local refinement. goCTF adopted such a cost-effective approach in estimating global focus gradient, without differentiating the tilt axis.

CTFTilt is a proven program specifically designed to determine global focus gradient assuming a flat sample within illuminated area. Local defocus can be extrapolated based upon modeled geometry as a single flat plane. CTFTilt performs an exhaustive search to determine tilt axis in the first step and subsequently estimates the tilt angle using conjugate gradient maximization of the calculated CTF and measured power spectrum. This is a computationally expensive approach to obtaining accurate and reliable results.

Alternatively, goCTF adopted a computationally inexpensive approach in estimating the global focus gradient, while still preserving adequate accuracy for the subsequent per-particle focus refinement. The tilt angle estimated by goCTF was observed to be consistent with the result produced by CTFTilt (Fig. S3). Interestingly, as explored in a previous study (Tan et al., 2017) using the benchmark dataset (EMPIAR-10097), local defocus extrapolated from the global focus gradient determined by CTFTilt, produced a slightly lower quality map compared with the gCTF focus refinement result, whereas goCTF has shown superior performance in comparison with gCTF. It would be worthwhile to explore the focus refinement performance impacted by the accuracy from a modeled global gradient in the future development of the goCTF program.

Recently, another software package, Warp, has been reported for real-time evaluation, correction and processing of cryo-EM data during their acquisition (Tegunov and Cramer, 2018). Warp has the capability to estimate local defocus without prior knowledge of a particle’s position. It divides a micrograph into patches by spatial grid control points for CTF estimation per patch. Subsequently, a tilted plane or a more complex geometry can be assessed and
modeled to recover local defocus at a given location. Warp presented a sound strategy in modeling the geometry, given an adequate signal obtained from each individual patch for CTF estimation. In practice, particularly in the case of a low-dose cryo-EM micrograph collected at high specimen tilt, focus gradient induced anisotropic scattering in combination with local drift could deteriorate a power spectrum significantly, thus limiting the accuracy of CTF estimation and impacting the model accuracy of the geometry plane.

As a benchmarking case (EMPIAR-10097), Warp reported a 3.9Å resolution reconstruction using software cryoSPARC with the original particle dataset CTF parameters replaced by Warp. In comparison, goCTF reported a 3.7Å resolution reconstruction following an identical approach, except with the original particle dataset CTF parameters determined by goCTF. This experiment suggests that both Warp and goCTF are equally capable of improving map reconstruction for single-particle cryo-EM tilt application.

The goCTF program inherits the same convention as CTFFIND4 with the input parameter values given interactively. The particle coordinate file corresponding to an input micrograph is required as a hidden input with a name suffix _go.star. The updated local defocus value is output in a separate file with a name suffix _goCTF.star.

## 5. Conclusions

The novel program goCTF was developed specifically to determine the CTF for single-particle cryo-EM customized for tilt applications. With its geometrically optimized approach, goCTF provides an ultra-reliable solution in per-particle focus refinement that is independent from the particle abundance at a given local region of interest in a micrograph. In addition, its iterative strategy for per-particle based focus refinement ensures accurate CTF determination that best represents a particle of interest without being overly constrained by the global focus gradient. Along with goCTF, the calculated global focus gradient plot and standard deviation defocus map, together with the heatmap of goodness fitting, could be adopted as additional means to evaluate the fitting performance prior to 3D refinement. The goCTF program has been validated and proven to improve final 3D reconstruction substantially in the near-atomic resolution range in the case of a published benchmark dataset with influenza hemagglutinin trimer.

In addition to increasing resolution of near-atomic range structures, goCTF also can improve our understanding of the structures of protein complexes at midrange resolutions. Particularly during the development of an experimental project with micrographs collected in suboptimal imaging condition, an intermediate resolution map is expected and equally important in leading to ultimate understanding of the structure of a protein complex, although it may suffer from preferred particle orientation. Apart from near-atomic resolution reconstructions, goCTF showed more potential compared to its peer program gCTF in preserving intermediate resolution with tilted datasets, due to its ultra-reliable performance in determining a global focus gradient for an image with extremely low SNR. Taken together, these qualities demonstrate that goCTF’s geometrically optimized approach enhances single-particle cryo-EM tilt applications, thus providing a reliable, effective and specimen-independent means to produce 3D reconstructions covering both the intermediate resolution range and atomic resolution.

The 4.1Å (gCTF) and 3.7Å (goCTF) resolution reconstruction volumes have been deposited into EMDB through access code EMD-8965 and EMD-8966, respectively. goCTF executable binaries are distributed under a XXX license and can be downloaded from: https://sites.google.com/umich.edu/min-su-ph-d/home.

## Acknowledgements

This work was supported by the Discretionary Fund of the University of Michigan Life Sciences Institute. I thank Dr. Melanie Ohi for many helpful discussions and for reading the manuscript. I thank Dr. Michael Cianfrocco, Dr. Jason Porta and Amanda Erwin for testing goCTF and providing their valuable feedback to further improve the program. I thank Christopher Lilienthal, Bradford Battey and Roy Bonser for support with scientific computing. I also thank Emily Kagey for proofreading the manuscript. I especially thank Dr. Amy Bondy for reading the manuscript and for suggestions in naming the program.

**Appendix A. Supplementary data**

